# Social benefits of facial experssion in a cichlid fish: Testing the face concentration hypothesis

**DOI:** 10.1101/2024.10.17.618800

**Authors:** Shun Satoh, Kazuya Fukuda, Hiroshi Matsui, Kento Kawasaka, Sayaka Matsuo, Takezo Banda, Kota Kanbe, Alu Konno, Shiro Takei, Masanori Kohda, Nobuyuki Kutsukake

**Author notes:** Equally contribution (Shun Satoh, Kazuya Fukuda).

## Abstract

The face is a uniquely distinctive stimulus, encapsulating a wealth of information. Among the myriad of social cues conveyed by the face, emotional signals, known as facial expressions, are paramount not only for humans but also for numerous social animals. The evolution of facial expressions in these animals can also manifest in taxa other than mammals, as suggested by various studies highlighting the socio-ecological benefits of facial expressions. In this study, we elucidated the social function of facial coloration, determined by melanophores, in the neotropical social cichlid *Symphysodon aequifasciatus*. In this species, facial coloration exhibits instantaneous changes in response to varying social contexts. Through behavioral observations and experimental manipulation, we confirmed that facial coloration in *S. aequifasciatus* serves to attenuate unnecessary aggressive competition among conspecifics. Furthermore, we observed that the facial area subjected to coloration in this species is innervated by the adenosine triphosphate- and noradrenaline-ergic nervous system. These findings indicated that facial expression in *S. aequifasciatus* depends on the sympathetic nervous system and has evolved independently of mammalian facial expressions. Our study highlights teleost fishes as valuable animal models for exploring the universality of facial expressions and their underlying cognitive mechanisms in vertebrates.

## Introduction

The modes of signaling in social interactions among animals are not solely determined by socio-ecological contexts, but can also be characterized in a taxon-dependent manner, such as in the case of bird calls ^1^. Within this broad spectrum of embodied social signals, which physical signals that serve as a ubiquitous medium for interaction across the animal kingdom, human faces are remarkably unique as they encapsulate a large amount of information and social cues related to various aspects, including sex ^2^, identity ^3^, emotional states ^4^, and attentive states ^5,6^. In particular, humans can manifest emotions through facial expressions—complex, socially communicative movements governed by specialized neural systems acting on facial muscles ^7,8,9^—that can, in turn, influence the behavior of others. It was Charles Darwin who initially recognized the significance of comparing the facial expressions of animals and humans, along with their associated cognitive abilities ^10,11,12,13^. Since that time, a numerous approaches for elucidating the evolutionary trajectory of facial expressions involves juxtaposing them with patterns of emotional expression observed in animals ^14^.

While the degree of comparability between human and animal facial expressions still remains contentious, recent research has provided evidence that intricate facial muscle movements in non-human primates function as facial expressions observed in humans ^14–16^. Moreover, even under a more restrictive definition of facial expressions—as social or emotional signals characterized by immediate changes conveyed through facial cues—certain animals such as laboratory mice^17^ and macaws^18^, also exhibit discernible facial expressions. The proximate mechanisms such as innervation of facial nerves and complexities of facial expressions in these species differ from those of primates. Nevertheless, these varieties of expressions share the utilization of facial structures to convey social signals. Based on this perspective, many studies report that many animals employ various facial organs to express social signals ^19^, while we may not generally recognize these as ‘facial expressions’. For example, displays of mouth and gill spreading, so-called frontal display, in many fishes may function as aggressive social signals^20^.

Interestingly, evidence from some mammals support that facial signaling functions as vehicles for conveying information and social cues related for example to identity, dominance rank, and sex ^14, 21^, alongside emotional signals. In these animals, multiple social cues are thought to be centralized in the face because of the evolutionary consequences of attention to the eyes ^14^. Moreover, studies in primates suggested that eyes serve as important signals to gaze direction and objects of interest by other individuals ^22,23^, resulting in social benefits for both signalers and signal receivers in social animals. Importantly, according to the body of evidence, we can predict that the centralization of social cues in the face, namely, a phenomenon wherein the significance of social signaling in face becomes more pronounced than that of body and multiple signals converge on the face, arises convergently in social animals rather than through lineage-based constraints.

Teleost fishes lack the elaborated facial musculature of mammals; therefore, it has been thought that facial expressions are absent in this taxon ^14^. However, recent studies have pointed to peculiarities in the faces of social fishes ^20, 24–34^. The eyes or face also appear to be an important area of focus for fish ^28, 32–34^. Kawasaka (2021) demonstrated that the eyes constitute the most critical feature characterizing the face in a particular group living cichlid *Neolamprologus pulcher* ^33^. As well as responding sensitively to eye, social fishes are known to initially focus on the face when presented with images depicting conspecifics ^28^, like primates do ^35^. The ubiquitous sensitivity of fish species to eyes and faces is further supported by the prevalence of eye-spot patterns on their bodies or fins as anti-predator coloration ^36^. Additionary, numerous species spanning various taxonomic groups have been documented to exhibit notable morphological or color variations localized to the face serving as social cues for inter-specific or individual recognition ^20, 24–29^. Moreover, it has been posited that fish possess specialized cognitive and neural processing mechanisms for facial recognition ^26, 30^ and that the facial area may also play a pivotal role in self-recognition in cleaner wrasse ^31^.

These investigations collectively underscore the face-related peculiarities in fish. In other words, we can safely say that social fishes, like primates, are sensitive to faces and eyes, utilize facial cues as central signals for functions such as individual recognition, and there is a possibility that possess specialized cognitive and neural mechanisms dedicated to perceiving these signals ^34^. Studying facial expressions in these animals, the vertebrates situated at the furthest phylogenetic distance from mammals (which have traditionally been the primary subjects of such studies), contributes to a deeper understanding of the general principles underlying the evolution of social signals emitted from face. However, to date, no studies have investigated facial signaling in fish, specifically, how dynamic changes in facial features influence the behavior of conspecifics.

To address this knowledge gap, we here focused on the social function of facial expression in the discus fish *Symphysodon aequifasciatus*, a neotropical cichlid species that dwells in the Amazon River basin ^37^. These fish reside in sunken trees alongside their conspecifics during the non-breeding dry season and reproduce in flooded forests throughout the breeding rainy season ^38^. Both male and female of reproductive pairs provide parental care and nourishment in the form of mucus for their offspring ^39^. Figure 1 illustrates the differences in body coloration of the discus fish and its related species from tribe Heroini (Banded cichlid *Heros severus*, triangle cichlid *Uaru amphiacanthoides*, flag cichlid *Mesonauta festivus*, and angelfish *Pterophyllum scalare*) when they exhibit aggressive behavior and when they are attacked by conspecifics (see supplemental information S1 for detailed observation methods). Discus fish, flag cichlids, and angelfish exhibit significant alterations in body coloration in response to social interactions among conspecifics, with these changes being notably more pronounced on the face than on the rest of the body (Fig. 1). Banded cichlids and triangle cichlids, on the other hand, social interactions among them is not accompanied by their body coloration changes and their faces do not show significant changes. Among these species, Satoh et al. (2016) demonstrated that discus fish can distinguish between their breeding partners and unfamiliar conspecifics based on the structural color patterns on their faces²_. We observed that the vertical black bar, composed of melanophores and situated above their eyes ^40^, undergoes marked alterations within ten seconds following the presentation of conspecific stimuli in discus fish (Supplemental Movie S2). Despite population and individual variations in the vertical black bars along the body flanks, facial black bars, so-called eye-patch pattern, are ubiquitous among discus fish_¹. Therefore, we selected the discus fish as our model species for studying facial expressions in fish.

**Figure 1.**
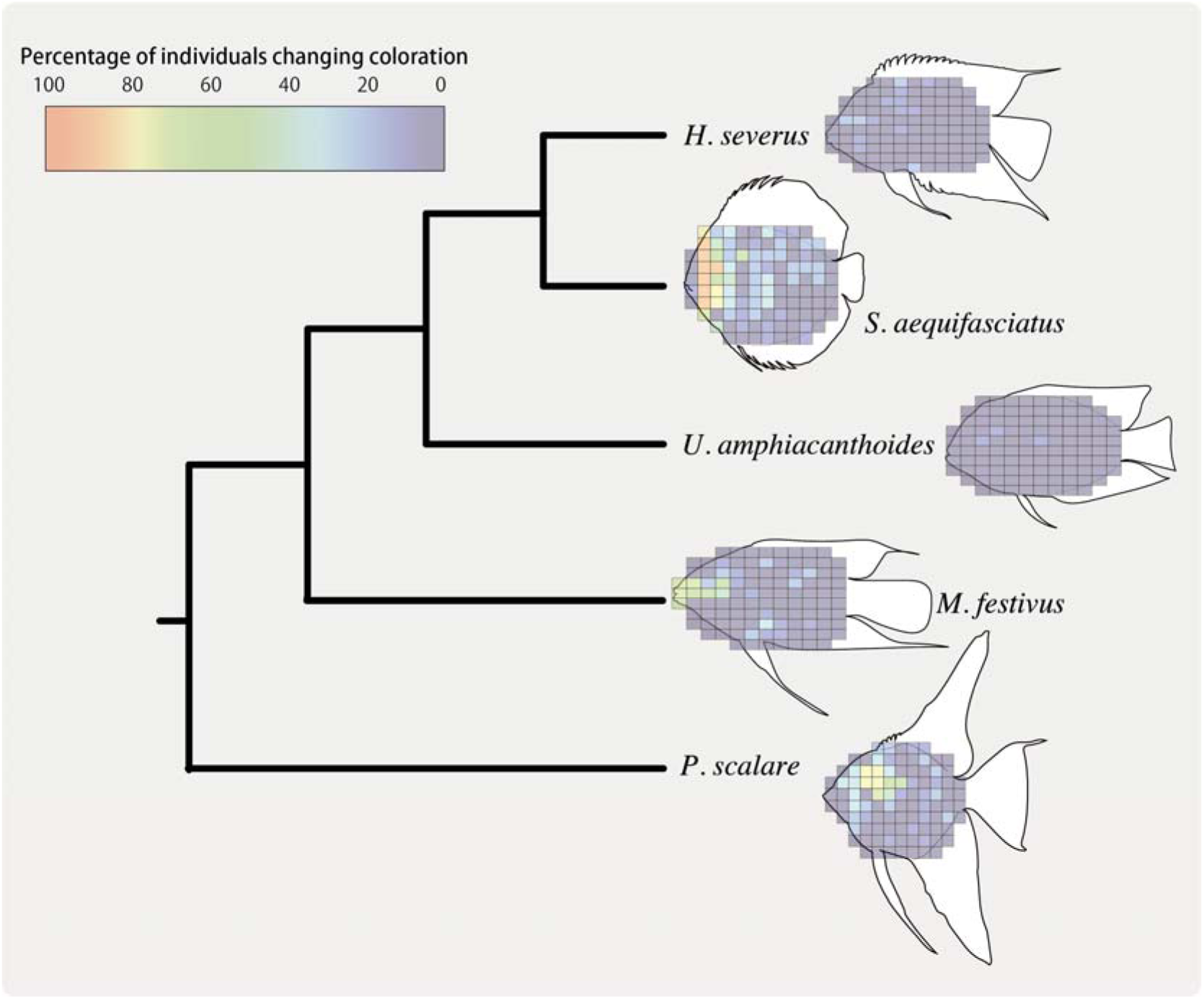
Differences in body coloration among discus fish and four related Heroini species when displaying aggressive behavior or experiencing conspecific aggression. The color coding indicates the percentage of individuals exhibiting pattern changes associated with these social contexts (aggressive behavior = 1; aggression received = 0). Refer to Supplemental information S1 for detailed observation methods.

In this study, we aimed to clarify whether facial coloration in discus fish functions as an social signal by using two complementary approaches: behavioral experiments and neuroscientific methods. In particular, we evaluated whether immediate changes in facial coloration can be referred to as facial expression by testing whether variation in face color suppressed unnecessary social costs. Through a combination of behavioral observations and experimental manipulation of facial coloration, we seeked to elucidate the role of this phenomenon as a social signal which benefits both signlar and reciver. Moreover, using experiments to determine which neurotransmitters activate melanophores in the facial area ^40^ and the slightly modified DEEP-Clear method ^42^, a clearing method for exploring the peripheral innervation of facial areas subjected to coloration, we attempted to identify which peripheral nerves are responsible for face color changes in discus fish and examined whether these are analogous to facial expressions in mammals.

## Results and Discussion

### Association between social behaviors and facial pigmentation patterns

To explore the association between distinct social behaviors and pigmentation patterns, we conducted behavioral observations of free-ranging *S. aequifasciatus* individuals (n = 8 males and 4 females) under simulated natural conditions. The pigmentation pattern, which encompasses melanophores in discus fishes, was classified into three categories: 1) complete absence of all vertical bars on the body, 2) manifestation of solely the eye-patch pattern, and 3) manifestation of all vertical bars, including the eye-patch pattern (Fig. 2). Specifically, when exhibiting aggressive behavior toward conspecifics, individuals typically abstained from displaying eye-patch patterns (Fig. 2a, b). Conversely, when under attack (Fig. 2c) or during periods of rest (Fig. 2d) and foraging (Fig. 2e), they did show them. Interestingly, eye patch patterns disappeared during sexual behavior as well as during aggression (Fig. 2f). With regard to the third pigmentation pattern, in which all vertical bars are manifested, individuals did not show it frequently, regardless of social behaviors (Fig. 2).

**Figure 2.**
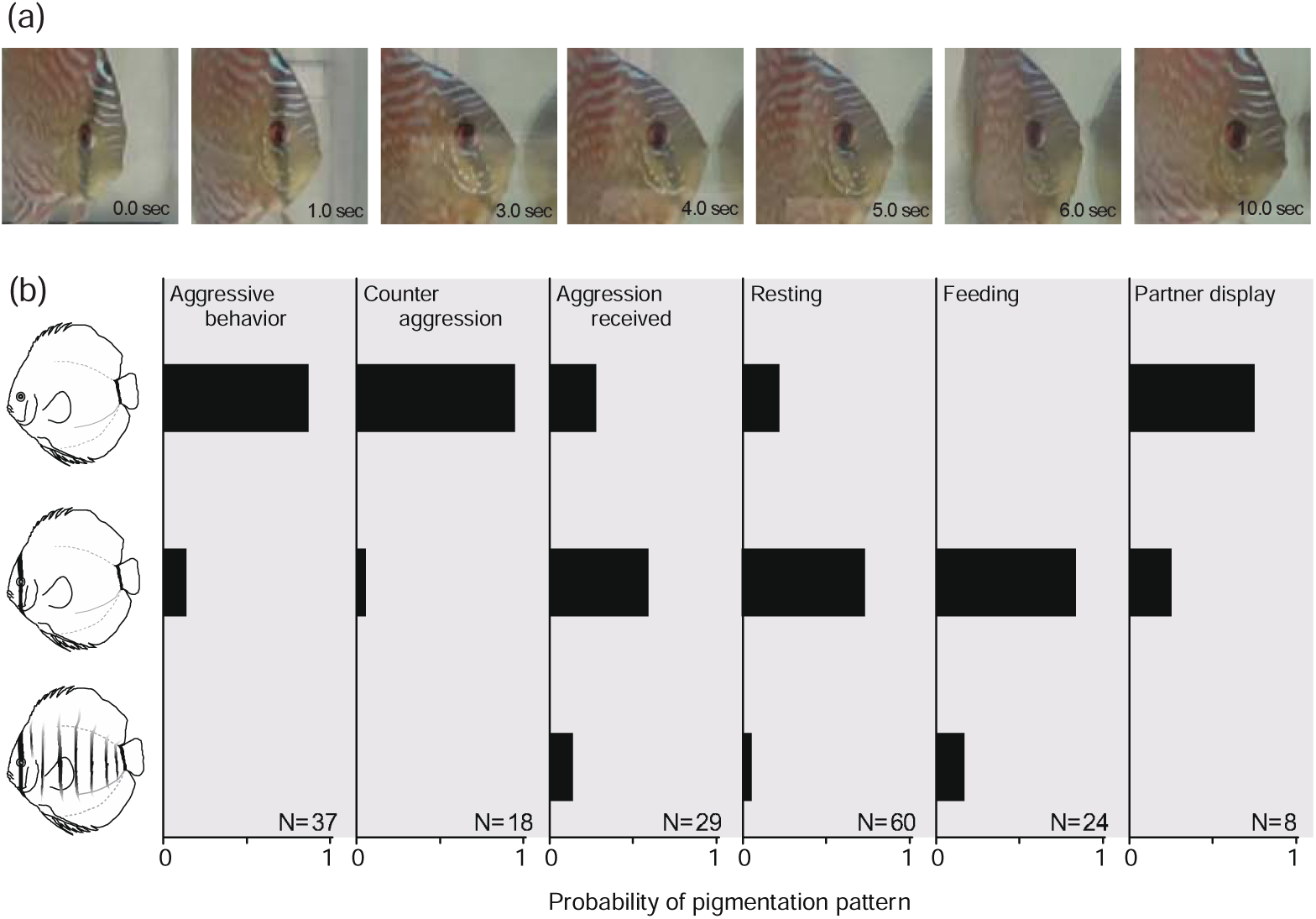
Facial coloration in discus fish *Symphysodon aequifasciatus*. Typical chronological changes of eye-patch pattern (a) and coloration associated with social behaviors (b). The bars represent the possibility of coloration patterns emerging when individuals engage in specific social behaviors. Three identifiable coloration patterns were detected: 1) absence of any vertical black bar composed of melanophores; 2) presence of only one bar consisting of melanophores, eye-patch pattern, and 3) presence of multiple bars throughout the body.

The pigmentation patterns in *S. aequifasciatus* were clearly correlated with certain social behaviors and were subject to rapid alteration, as shown in Supplementary movie S2. Numerous instances of body color change serving as a social signal have been documented in various social fishes. For example, in the group-living cichlid *Neolamprologus pulche*r ^43^ and mouth-brooding cichlid *Astatotilapia burtoni* ^40,44^, the intricate pigmentation pattern in the facial area serves as a social signal indicating aggressiveness, as similarly observed in discus fishes. However, there is no conclusive evidence that these changes in pigmentation pattern occur immediately in conjunction with the specific display, which would reflect a motive for aggression associated with social rank over a longer term, as suggested by Rodrigues et al. (2009) ^45^. Color change in the facial area in discus fishes is more akin to facial expression reported in primates ^19^ than to variation in facial coloration reported in other fish species.

The expression of distinct emotional states on the human face is characterized by the subtle movements of facial musculature ^7–9^. Conversely, in discus fishes, it is the alteration of pigmentation patterns in the facial area generated by the contraction of chromatophores, rather than facial muscles, that leads to the manifestation of specific facial expressions that correspond to specific emotional states. In particular, in this study, *S. aequifasciatus* individuals showed the eye-patch pattern when they were being fed or had received aggression but did not display it when engaged in aggressive or partner-specific behaviors. This suggested that the presence of the eye-patch pattern may be indicative of normalcy (submissive behavior) while its absence may denote excitement (aggressive behavior).

### Benefits of facial coloration patterns as social signals

To examine whether specific facial coloration patterns influence the degree of same-sex sexual harassment perpetrated by dominant individuals, we observed a series of aggressive behaviors between subordinate and dominant individuals for 20 minutes. We specifically focused on whether a subordinate that was under attack could decrease the probability of being subjected to continuous harassment (defined as an attack occurring within 5 seconds of the first attack) from a dominant individual by modifying its eye-patch pattern.

When a subordinate individual was attacked, alterations in facial coloration significantly affected the occurrence of persistent harassment (binomial generalized linear mixed model [GLMM], df = 3, χ^2^ = 63.589, p < 0.001) (Fig. 3). Subordinate individuals who did not exhibit the eye-patch pattern and did not modify their facial coloration upon being attacked by dominant individuals were significantly more likely to experience continuous harassment compared with those who did (df = 1, χ^2^ = 54.186, p < 0.001). Similarly, in subordinate individuals who maintained the eye-patch pattern before and during harassment, the possibility of continuous attacks also decreased (df = 1, χ^2^ = 10.05, p = 0.002). Conversely, the likelihood of continuous harassment remained comparable when subordinate individuals without the eye-patch pattern failed to alter their facial coloration during aggression, regardless of whether those initially displaying the pattern modified their facial coloration (df = 1, χ^2^ = 2.093, p = 0.148). Our behavioral observations suggested that alterations in facial coloration serve to deter harassment from dominant individuals in *S. aequifasciatus*.

**Figure 3.**
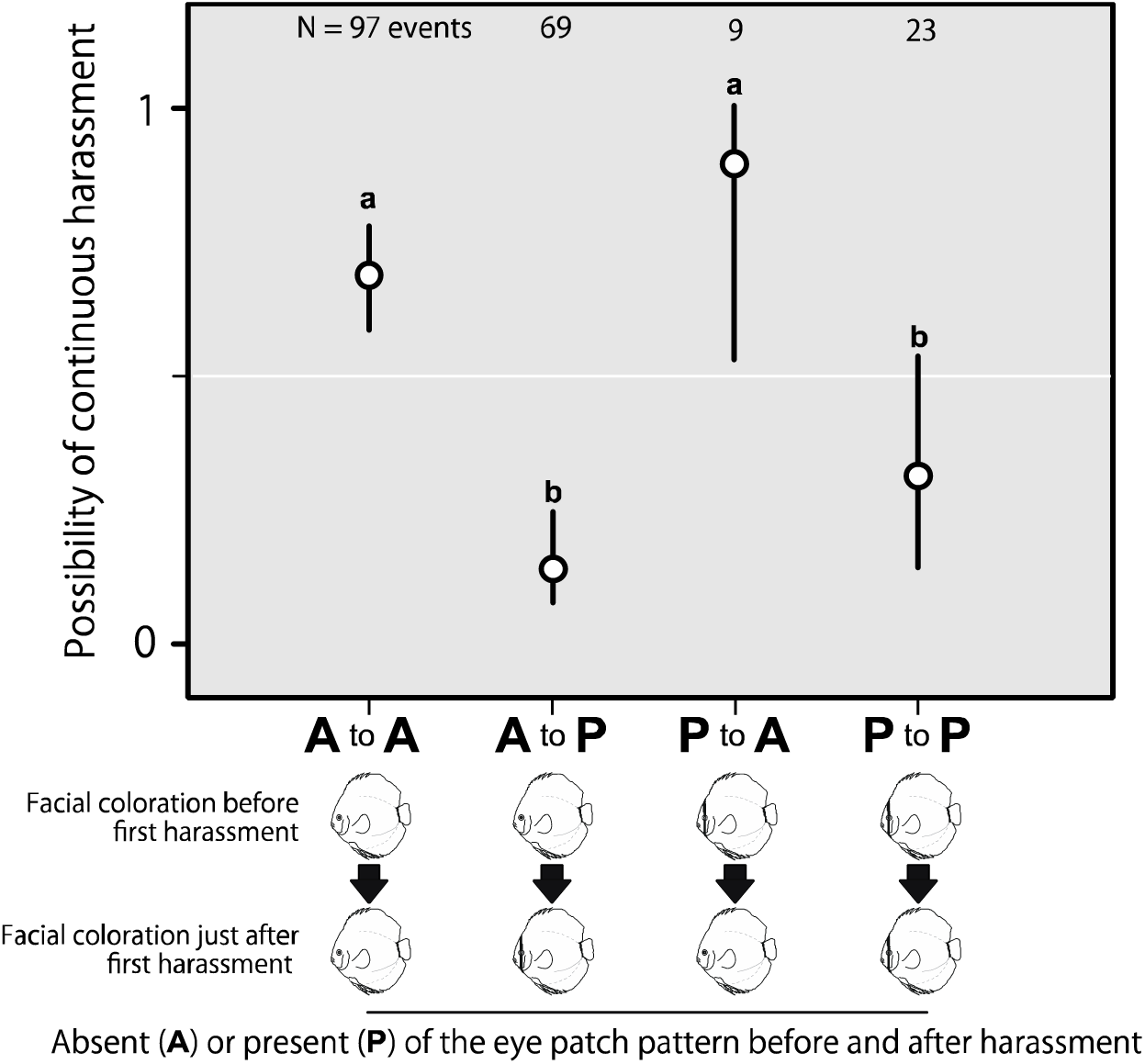
Effects of facial coloration in discus fish *S. aequifasciatus* subordinates on the continuous aggressive harassment by dominant individuals. The data are presented as mean values along with their corresponding 95% confidence intervals, and the sample sizes are indicated above the bars. Significant differences were determined using post-hoc binomial GLMMs, with p-values adjusted using Bonferroni correction (α level = 0.0083). The possibility of experiencing continuous harassment was notably higher in *A to A* than in *A to P* (df = 1, χ^2^ = 54.186, p < 0.001) and *P to P* (df = 1, χ^2^ = 10.05, p = 0.002), but similar to that of *P to A* (df = 1, χ^2^ = 2.093, p = 0.148). Conversely, the probability of continuous harassment was significantly lower in *A to P* than in *P to A* (df = 1, χ^2^ = 22.076, p < 0.001), but similar to that of *P to P* (df = 1, χ^2^ = 3.3123, p = 0.0688). Additionally, there was a significant difference in the possibility of continuous harassment between *P to A* and *P to P* (df = 1, χ^2^ = 9.690, p = 0.002).

However, because facial coloration in discus fishes may be correlated with a variety of social behaviors and psychological motivations ^20^, it is difficult to determine whether it has a specific function in inhibiting harassment from dominant individuals. To overcome this limitation, we conducted experiments to manipulate facial coloration in *S. aequifasciatus* using rearing method based on light exposure commonly employed by ornamental fish keepers (see methods). As shown in several studies conducted in the field of aquaculture science ^46^, exposing juvenile fish to intense light can affect the melanophore-derived body coloration. Therefore, we obtained *S. aequifasciatus* individuals in which the function of melanophores was abolished by exposing them to intense light during the juvenile stage. The manipulated discus fish were unable to exhibit melanophore patterns regardless of social context (Supplemental figure S3), and thus were considered to be unable to convey proper social communication signals. In the behavioral experiment, we observed changes in aggressive interaction over time in experimental pairs in which 1) both individuals could properly display the eye-patch pattern (WW pairs, n = 8 pairs), 2) one individual was light-treated (WL pairs, n = 6 pairs), and 3) both individuals were light-treated (LL pairs, n = 7 pairs).

We detected a significant two-way interaction between the first day of the behavioral experiment and the type of experimental pair in terms of the frequency of aggressive interactions over a period of 20 minutes (negative binomial GLMM, df = 2, ^2^ = 67.968, p < 0.001). In the WW group, the frequency of aggressive interactions between same-sex individuals decreased day by day, but in the LL group, which was composed of pairs in which both individuals who could not express optimal social signals, the frequency of aggressive interactions did not decrease within days (Fig. 4). In the LW group, the variation in the frequency of aggressive interaction was in-between the values observed for the WW and LL groups (Fig. 4). In this experiment, the manipulation inevitably affected not only facial coloration but also melanophore-derived pigmentation patterns in the rest of the body. However, considering that pigmentation patterns other than those in the face were largely unchanged in social displays between conspecifics (Fig. 2), the obtained results can be safely interpreted as being caused by the inability of facial melanophores to produce patterns, which therefore prevented proper communication between conspecifics.

**Figure 4.**
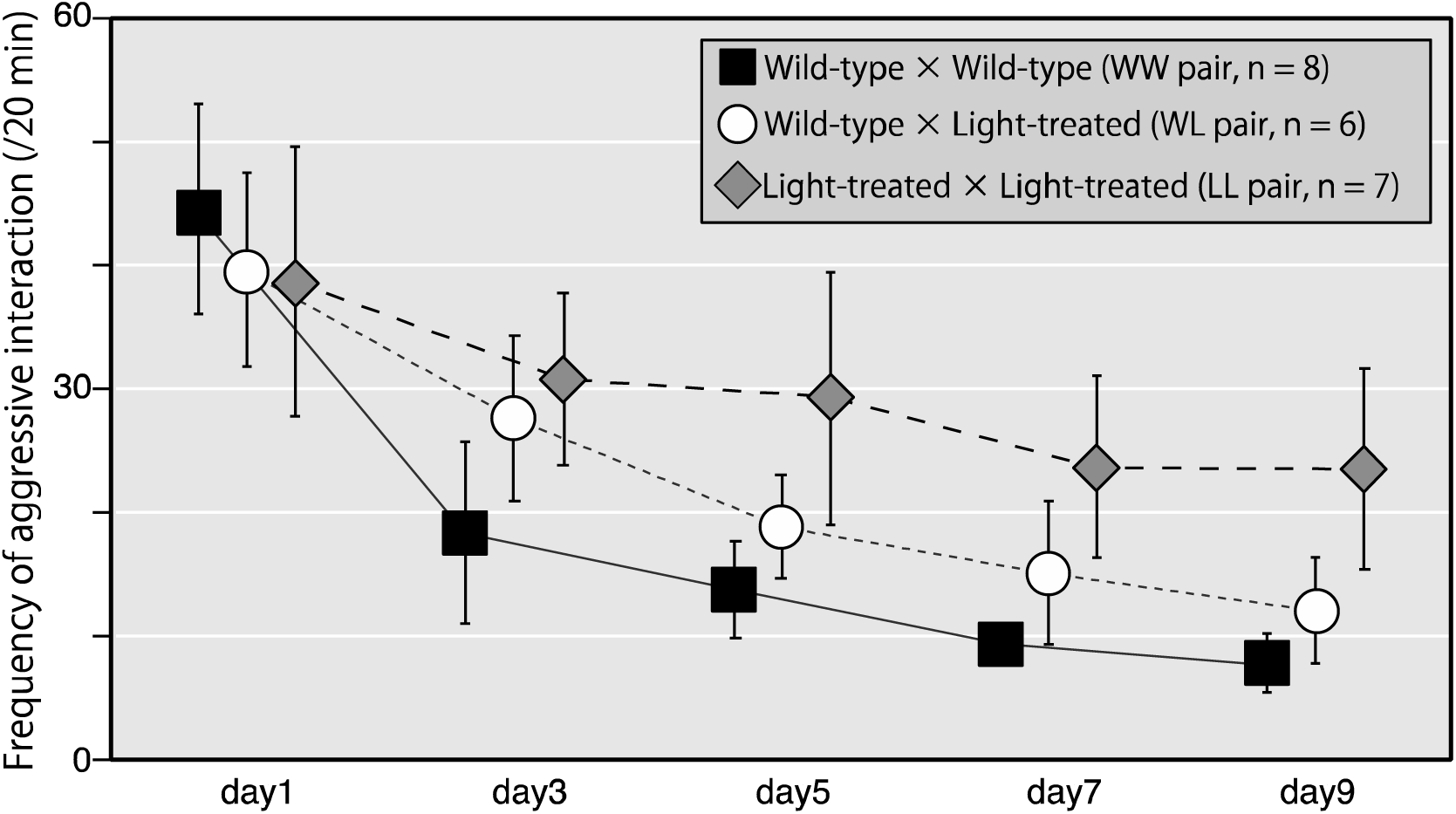
Effects of the experimental manipulation of melanophores on aggressive social interaction in discus fish *S. aequifasciatus*. The figure shows the relationship between the frequency of aggressive interaction within experimental groups and days since the start of the experiment. The experimental groups, consisting of wild-type × wild-type individuals (WW), wild-type × light-treated individuals (WL), and light-treated × light-treated individuals (LL), are indicated by black, white, and gray bars, respectively. A significant interaction was detected between the type of experimental group and the duration of the experiment (negative binomial GLMM, df = 2, ^2^ = 67.896, p < 0.001). Data are presented as mean values ± standard deviation (SD).

During the experimental days, individuals lost 0.5%–8.4% of their body weight. The percentage of the reduction in body weight significantly varied among experimental groups, with LL suffering the greatest weight losses, followed by WL and WW, in that order (linear mixed models [LMM], df = 2, F = 4.398, p = 0.025) (Fig. 5). Moreover, whether individuals became dominant (or subordinate) during the experiment also had a marginal effect on body weight reduction (df = 1, F = 3.955, p = 0.061). Sex (df = 1, F = 2.354, p = 0.145) and whether individuals had been subjected to the light treatment (df = 1, F = 0.224, p = 0.641) had no significant effects on the reduction. Social signals have been shown to confer benefits to both conveyors and recipients ^47^. The differential reduction in body weight observed across experimental groups in this study, irrespective of social hierarchy, aligns with the tenets of classical adaptive signaling theory and provides compelling evidence that facial coloration yields socio-ecological benefits in *S. aequifasciatus*. This species forms social aggregates in the Amazon River, an environment characterized by high predation pressure ^38^, and the act of leaving the group, even to assume a subordinate position, carries fitness costs ^48^. Here, the individuals whose ability to modify facial coloration had been manipulated were possibly unable to express the submissive signal toward dominant individuals and thus establish a dominance hierarchy. Facial coloration may serve to mitigate superfluous competition within social aggregates characterized by dominance hierarchies.

**Figure 5.**
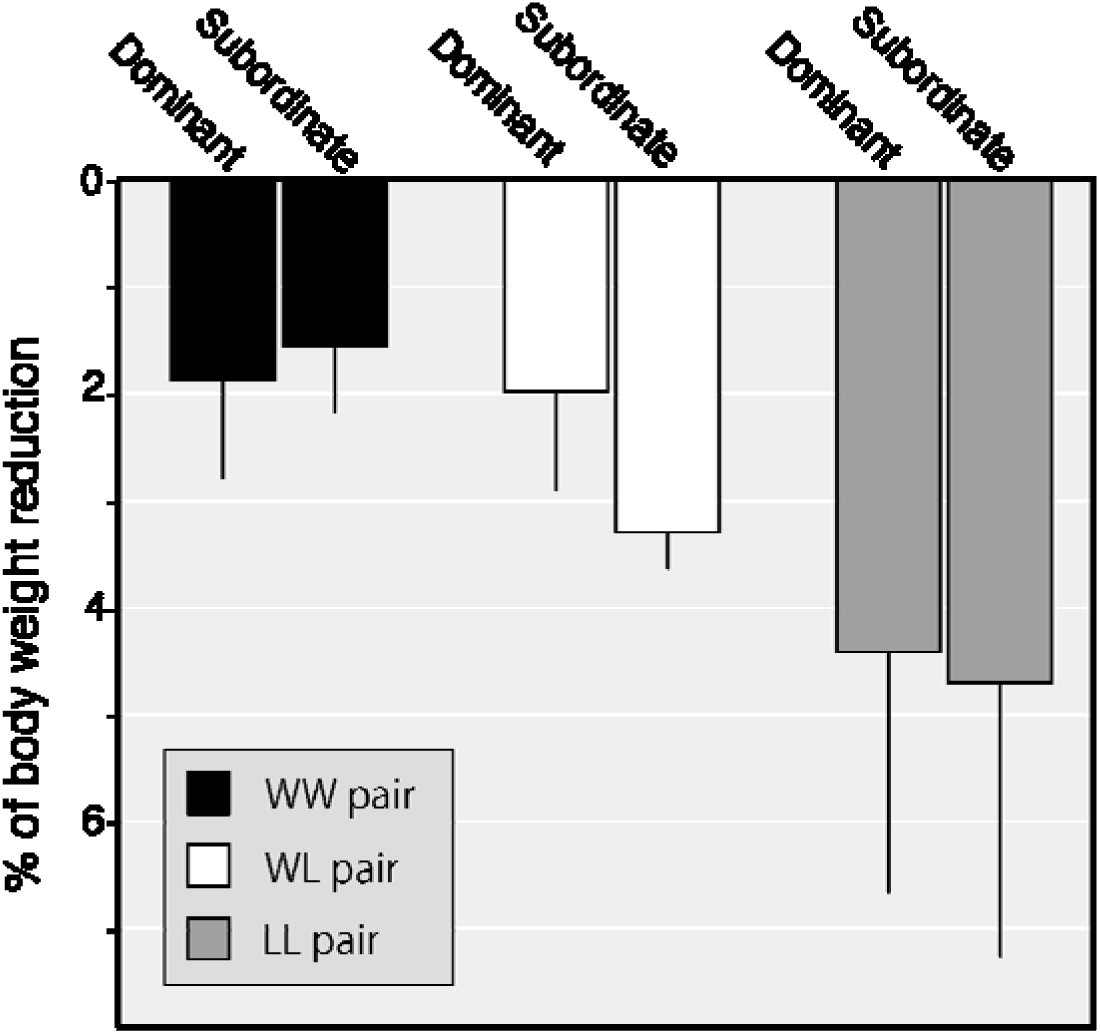
Reduction in body weight during the 9-day experiment in dominant and subordinate discus fish *S. aequifasciatus*. The experimental groups, consisting of wild-type × wild-type individuals (WW), wild-type × light-treated individuals (WL), and light-treated × light-treated individuals (LL), are indicated by black, white, and gray bars, respectively. Significant differences were detected in the percentage of reduction in body weight between experimental groups (LMM, df = 2, F = 4.398, p = 0.025), while marginal differences were observed in dominance rank (df = 1, F = 3.955, p = 0.061). Data are presented as mean values ± standard deviation (SD).

### Peripheral innervation regulating facial coloration

The results of our behavioral experiments indicated that facial coloration in *S. aequifasciatus* functions as a social signal in a similar manner to human facial expressions. One approach to confirm whether the two phenomena are indeed analogous is to compare their underlying regulatory mechanisms. Therefore, we conducted two experiments to elucidate the types of peripheral nerves innervating the facial area subjected to changes in coloration in *S. aequifasciatus*. Generally, discus fishes exhibit two primary mechanisms of color change: physiological color change, mediated by hormonal and neural regulation of peripheral nerves, and morphological color change, involving the accumulation or depletion of chromophores ^49–51^. Previous behavioral experiments have shown that changes in facial coloration in discus fishes occur rapidly, within a few seconds. Since morphological color changes usually require days to weeks ^51^, we hypothesized that the above-mentioned rapid changes are physiological, which are known to occur over shorter periods. In particular, they may be under neural control, which has been shown to regulate very rapid color changes within seconds ^44, 49^. In discus fishes, body color changes, particularly those involving black coloration via the aggregation and dispersion of melanosomes within melanophores, are generally regulated by the sympathetic nervous system ^44, 52, 53^. Sympathetic postganglionic neurons in discus fishes are known to possess several neurotransmitters, including noradrenaline (NA), adenosine triphosphate (ATP), neuropeptide Y, and acetylcholine ^50, 54^, the first two of which are the primary factors influencing the motile activity of melanosomes ^49, 55^. Therefore, we first examined whether NA and ATP control facial coloration in discus fish by administering these neurotransmitters ex vivo. In this experiment, scales (n = 20) with chromatophores were freshly collected from eye patches and observed under a microscope. The scales were then immersed in neurotransmitters (1 µM NA or 10 mM ATP) or in PBS, which was used as a control. We quantified the rate of change in melanosome dispersion (%) in the scales and observed significant differences between ATP exposure and PBS exposure (two-way interaction between time and treatment, df = 1, F = 200.888, p < 0.001 by LMM), with melanosomes showing dispersion in the ATP exposure group (Fig. 6a, c). Conversely, NA exposure resulted in significant melanosome aggregation compared to PBS exposure (two-way interaction between time and treatment, df = 1, F = 89.651 p < 0.001) (Fig. 6b, d). These results suggested that changes in facial coloration in discus fish also depended on sympathetic innervation. To further validate this hypothesis, we investigated whether sympathetic postganglionic neurons (noradrenergic neurons) reached the eye-patch region via tissue clearing. Tissue clearing is a methodology to visualize appropriately labeled structures in chemically cleared, three-dimensional tissues, which is especially useful to track neurons or blood vessels ^56, 57^. We cleared the heads of X individuals using the slightly modified DEEP-Clear method ^42^ and performed immunohistochemical staining using an anti-tyrosine hydroxylase (TH) antibody (the key enzyme that converts tyrosine to L-DOPA, the initial step in NA biosynthesis). Finally, we performed refractive index matching using CUBIC-R reagent ^58^. The results showed that TH-immunoreactive (TH-ir) fibers reached the eye-patch region (Fig. 6e). Furthermore, morphological observations indicated that these fibers connected to the scales containing the melanophores constituting the eye patch (Fig. 6f, g). These findings strongly suggested that sympathetic nerves innervate the facial area subjected to changes in coloration in discus fish.

**Figure 6.**
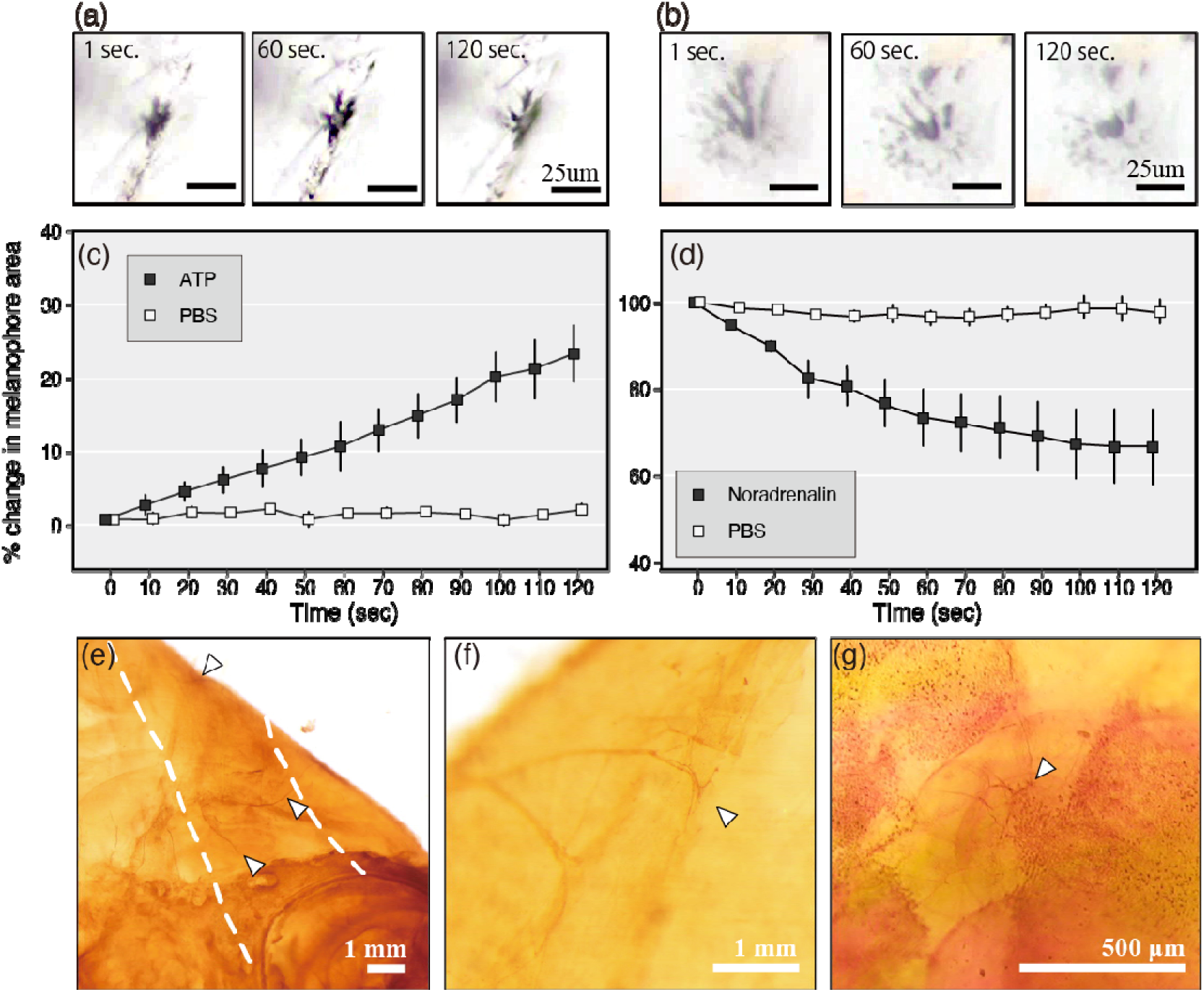
Effect of the noradrenaline (NA) and adenosine triphosphate (ATP) neurotransmitters used by the sympathetic postganglionic neurons on melanophores constituting the eye patch and tyrosine hydroxylase-immunoreactive projection to the eye-patch region in discus fish *S. aequifasciatus*. The peripheral nerves regulating the melanophores constituting the eye patch were examined. **a–d** Temporal variations in the extent of melanosome dispersion upon the exposure of melanophores constituting the eye patch to NA and ATP . The rate of variation in melanosome dispersion upon exposure to ATP (a, c) and NA (b, d) was significantly different from that observed in the PBS control (two-way interaction between time and treatment, both p < 0.001). Data are presented as mean values ± standard error. **e–g** The tissues were cleared and immunostained using an antibody against tyrosine hydroxylase (TH), revealing that TH-immunoreactive (TH-ir) projections extended to the eye-patch region (white arrowheads). The white dotted lines indicate the eye-patch region (**e**). The observations strongly suggested that TH-ir fibers terminate at the scales possessing melanophores forming the eye patch: (**b**) specimen whose scales had been removed to improve the visibility of the TH-ir fibers and **(e)** specimen with intact scales.

Our results suggested that, unlike human facial expressions, which are regulated by facial nerves ^7^ via the neurotransmitter acetylcholine ^59^, facial coloration in the examined discus fish is regulated by the sympathetic nervous system. Facial coloration serves as a social signal and plays a significant role in social interactions with other individuals ^60, 61^. Therefore, its functioning to convey facial expression and its peripheral neural regulation observed in this study may not be limited to discus fish but could be prevalent among social fish species. One of the interesting aspects of our findings is that, while facial expressions and coloration similarly function as social signals in humans and fish respectively, they are controlled by different nervous systems. In humans, they are regulated by voluntarily as well as in involuntarily through facial nerves, whereas in fish they are controlled by the autonomic nervous system.

Regulation by voluntary nerves implies that humans can deceive others by displaying facial expressions that do not reflect their authentic emotions ^62.63^. Recently, it has been reported that fish also possess the ability to deceive others tactically ^64^. However, considering that facial coloration in fish is regulated by the autonomic nervous system, it is unlikely that they can use neurally controlled rapid variations in facial color for intentional deception. Although it is beyond the scope of this study to determine whether fish facial expressions are intentional or volitional, facial expressions of discus fish are likely a direct manifestation of their emotional states.

### Evolution of facial expression in social fishes

Of the body coloration composed of melanophores, discus fish intensively linked and changed only facial coloration upon engaging in social behavior toward conspecifics. Our findings further validated that facial coloration in this species serves to mitigate superfluous aggressive competition among conspecifics, thereby benefiting both signalers and recipients within the dominance hierarchy. This supports the hypothesis that facial coloration in discus fishes exhibits functional parallels to mammalian facial expressions, predominantly those observed in primates^19^. Conversely, the neural mechanisms underlying facial coloration in these fishes markedly differ from those regulating facial expressions in mammals, strongly suggesting a convergent evolution of facial expressions in each lineage.

In humans, facial expressions are hypothesized to have initially evolved as a mechanism for modulating physiological states within an individual (e.g., widening of the eyes in expressions of fear, facilitating enhanced peripheral vision) before assuming a secondary role as social signals ^67^. The vertical bars present on discus fishes, which are typically found among driftwood, may serve as forms of cryptic or disruptive coloration ^68^. Indeed, when confronted with predators or subjected to prolonged intra-species aggression, discus fishes display vertical bars, including eye-patch patterns, spanning their entire body ^41^. Similarly to facial expression in humans, facial coloration in discus fishes may have initially evolved as a cryptic form of coloration mediated by melanophores, before subsequently being repurposed as a means to convey submissive social signals localized solely to the facial region, undergoing coevolutionary adaptations to facilitate their perception. Moreover, as an alternative prospective evolutionary scenario other than cryptic coloration, the signals linked to long-term hierarchical dominance cues within the facial region ^40,43,44^ may have undergone evolutionary divergence toward immediate modification in response to social stimuli.

When a fish is threatened by predators, triggering a fight-or-flight response analogous to harassment by conspecifics, it experiences acute stress and often exhibits cryptic body coloration^69,70^. This stress activates the sympathetic nervous system, rapidly increases systemic cortisol levels, and raises plasma glucose levels, creating a physiological state conducive to fight-or-flight responses against the stressor ^69–71^. A similar situation assumably occurs when subordinate individuals try to avoid harassment from dominant individuals ^69,72–74^. Therefore, it should be plausible to hypothesize that the mechanism regulating variations in body and facial coloration via the sympathetic nervous system to escape from predator-related threats has secondarily adapted to cope with similar behavioral and physiological contexts (for example to avoid harassment from dominant individuals), even though the stressors are different.

Importantly, facial coloration in discus fishes changes independently from body coloration (Fig. 2), suggesting the presence of distinct control mechanisms for these areas. It is conceivable that discus fishes have developed neural mechanisms to gradually or alternatively change the coloration of their face and body in response to different stimuli and social contexts. Alternatively, differentiated neural pathways controlling facial and body coloration may have evolved, as exemplified by the unique peripheral nerve branches found in these fish species, such as the recurrent facial taste neurons in sea catfish, which have acquired taste buds on their body surface and fins in response to socio-ecological demands over the course of evolution ^75^.

The hypothesis that signals are concentrated in the face is further supported by evidence from other socio-cognitive functions, including individual recognition ^25,27–31^, self-recognition ^33^, attention^30^, and specialized cognitive processes in fish ^28,32^.To the best of our knowledge, there have been no reports documenting instantaneous social signals on the faces of fish, as observed in discus fishes. However, this may stem from a paucity of research on facial expressions in this animal group rather than from an absence of such expressions. Indeed, in numerous fish species, social signals indicative of long-term dominance hierarchies are localized to the facial region ^40,44,45^, despite the face being an anatomically limited part of the body. We verified that the immediate (specifically, about 10 seconds: Fig. 2a) change in discus facial coloration functions as a social signal, and that this signal may be an emotional expression for this fish. This finding suggests that emotional social signals are likewise concentrated in the face of fish where the importance of the face in communication has recently been noted (Kohda et al. 2024).

A comparative study conducted on primates has revealed a correlation between the efficacy of facial expression and social complexity ^76^. Comparative analysis which fish species have developed facial expressions would hence be the next step to future study. Indeed, related species of discus fish frequently change their facial patterns during social interactions, while others do not (Fig. 1). The present study, by following a similar line of research and exploring the relationship between social systems and facial coloration in a discus fish, contributes to comprehensively elucidating the evolution of facial expression within this taxon. It also demonstrated that teleost fishes are valid animal models for probing the generality of facial expressions and their underlying cognitive mechanisms in vertebrates. An integrated approach to the function, complexity, and mechanisms of signaling of embodied communication in face will be needed to fully understand the general rules of evolution in facial expersison in social animals.

## Methods

### Subjects and fish husbandry

Mature male and female *S. aequifasciatus* specimens were procured from ornamental fish traders that imported them from the Brasilia Amazon and bred F1 individuals in Japan. These specimens were housed in four separate water tanks. Uniform fish density and sex ratio were meticulously maintained across all stacked tanks. Water temperature and pH were carefully regulated and kept within the ranges of 27.0°C–30.0°C and 5.0–7.0, respectively. The fish were nourished with an artificial diet formulated from beef heart and mosquito larvae (three times per day). The sex of specimens was determined based on their distinctive body coloration and morphological characteristics. In total, eight males and four females were used for *behavioral observation I*, and 20 males for *behavioral observation II*. All observations and experiments were conducted from 12.00 to 15.00 and recorded using a video camera (HDR-CX470 and HDR-CX370, Sony Corporation, Tokyo, Japan) placed at ca. 30 cm in front of the water tanks.

### Behavioral observation I: pigmentation patterns of S. aequifasciatus in different social contexts

The behavior of *S. aequifasciatus* in a water tank was observed to determine what social context correlated with pigmentation patterns, which change instantly (Supplemental movie S2). Driftwood and dry leaves were placed in the tank (360 L, 120 × 60 × 50 cm), and the water was colored using humic acid to mimic the natural habitat ^36,41^. Three mature individuals (two males and one female) were held in this tank for 3 days to allow accustomization to the environment. During behavioral observations, which lasted for 25 min, mosquito larvae were fed to the fish to promote social interaction among them, and the pigmentation patterns of focal fish were evaluated when they showed six specific social behaviors (i.e., aggressive behavior, counter aggression, received aggression, feeding, and partner display) based on the ethogram described in Satoh et al. (2016)^25^. Aggressive behavior was defined as a fish approaching the lateral side of a non-partner while opening its mouth and spreading its gill covers and fins as well as a fish aggressively biting and/or bumping into the non-partner. A focal fish receiving aggressive behaviors from other experimental fish and then showing the same behaviors was defined as counter-aggression, while the lack of any behaviors upon receiving aggression was defined as received aggression. Partner display was defined as two fish approaching each other holding up their heads and folding their pelvic fins toward their body. This type of behavior is observed when fish form monogamous reproductive pairs ^25^. Additionally, rest was defined as a state in which the above behaviors were not exhibited; specifically, pigmentation patterns in this state were recorded about 1 minute after the other behaviors were observed. Observations of the rest state were conducted randomly five times in eight male and four female adults.

Three types of pigmentation patterns were detected in *S. aequifasciatus* (Fig. 2). Typically, discus fishes have a total of eight vertical bars on their body, including the facial area ^41^. However, in this study, individuals did not show vertical bars on their body flank when such bars were absent on their facial area.

### Behavioral observation II: effects of facial coloration on aggressive behavior

The objective of these observations was to investigate the impact of facial coloration on the frequency of harassment by dominant individuals toward subordinates. To this end, 10 pairs of mature males (n = 20) were placed within an experimental water tank (162 L, 60 × 45 × 60 cm). In these same-sex experimental pairs, the fish differed in body size by a minimum of 1.5 cm, which ensured the establishment of a size-based hierarchy between them; smaller individuals were designated as subordinates, while larger individuals assumed the role of dominants. The experiment was focused on determining whether the presence of an eye-patch pattern in subordinate individuals served as an social signal to mitigate the possibility of continuous aggression (harassment) from dominant individuals, given that eye-patch patterns had been frequently displayed by subordinates in response to aggressive behavior from conspecifics (Fig. 2).

First of all, upon observing aggressive behavior by dominants against subordinates, the facial coloration of the latter (i.e., absence or presence of the eye-patch pattern) was recorded (hereafter called ‘first aggression’). After the first aggression, whether the dominant individuals continuously attacked the subordinates was also recorded. In this study, an aggressive behavior that occurred within 5 seconds of the first aggression was defined as continuous harassment. If this situation was produced, the facial coloration of subordinates was recorded. In contrast, if dominant individuals did not continuously attack, the facial coloration of subordinates was recorded 5 seconds after the first aggression. If dominant individuals repeated the aggressive behavior three or more times within a single sequence of aggressive behaviors, these were considered as part of the first aggression, and observations continued in the same manner until the continuous harassment ended. During these observations, which lasted for 20 min, a maximum of six consecutive aggressive behaviors against subordinates occurred within a single behavioral sequence. Instances where subordinates did not exhibit the eye-patch pattern upon first aggression by dominants and subsequently maintained their facial coloration without alteration (*A to A*) were observed 97 times. Instances where subordinates did not exhibit the eye-patch pattern upon first aggression but did so after it (*A to P*) were observed 69 times. Instances where subordinates exhibited the eye-patch pattern upon first aggression and subsequently altered their facial coloration to a non-eye-patch pattern (*P to A*) were observed 9 times. Finally, instances where subordinates exhibited the eye-patch pattern upon first aggression and subsequently maintained their facial coloration without alteration were observed 23 times. It should be noted that in 20 cases of aggression, facial coloration could not be discerned; therefore, these data were removed from subsequent analyses

### Experimental manipulation of social signals

To investigate the potential function of facial coloration as a social signal, body coloration in *S. aequifasciatus* was experimentally manipulated using a light treatment known among discus fish enthusiasts that targets the vertical bars, which are believed to serve as social signals. Specifically, this treatment aimed to eliminate the vertical bars by exposing juvenile fish to intense illumination. Initially, mature females and males exhibiting partner displays (Satoh et al. 2016) were transferred to separate water tanks (122 L, 60 × 45 × 45 cm or 91 L, 45 × 45 × 45 cm). Substrate bricks suitable for spawning were provided in these tanks, and humic acid was periodically added to regulate water quality. Upon spawning, the offspring remained with their parents for 10 days, as early free-swimming fry rely on parental mucus for sustenance (Satoh et al. 2017). Subsequently, 50 fry from the same brood were placed in two smaller tanks. One tank received illumination from a light source positioned approximately 30 cm above, thereby constituting the light-treated group, while the other tank remained under ambient room lighting, serving as the control for wild-type individuals. The juvenile fish were fed *Artemia* spp. nauplii during the experiment. The lighting regime for the light-treated group was synchronized with the room lighting, maintaining a 12-hour light / 12-hour dark cycle. After 3 months, the juveniles were identified and moved to the same water tanks (162 L, 60 × 45 × 60 cm or 182 L, 90 × 45 × 45 cm) until they reached sexual maturity, which typically occurs over a period of approximately 1.5–2.0 years. Facial coloration was evaluated in the light-treated individuals in the same manner as in *behavioral observation I*, but none of them exhibited an eye-patch pattern (Supplemental figure S3).

Light-treated and wild-type individuals were used to test whether facial coloration functions analogously to emotional expression. Three types of experimental pairs were created: a group consisting of wild-type individuals only (WW pair, n = 4 male pairs and 4 female pairs), a group of wild-type and light-treated individuals (WL pair, n = 3 male pairs and 3 female pairs), and a group of light-treated individuals only (LL pair, n = 3 male pairs and 4 female pairs). These same-sex experimental pairs (42 individuals in total) were selected to ensure homogeneity in kinship and were allowed to meet each other for the first time. After measuring their body weight in 1 mg, they were placed in a water tank (122 L, 60 × 45 × 45 cm) for 9 days. During the experiment, the fish were individually fed with five mosquito larvae per day using tweezers. This was to ensure that the dominant individuals did not monopolize the food. The number of aggressive interactions between pair partners over a period of 20 minutes was counted on days 1, 3, 5, 7, and 9. In this study, aggressive interaction was defined as total aggressive display and direct aggression occurring between pair partners. After 9 days of observations, the body weights of the experimental pairs were measured again. Among the examined groups, individuals exhibiting more aggressive behaviors toward their experimental partners during the experiment were defined as dominant, while those receiving more attacks as submissive. The frequency of aggressive interaction was predicted to be higher in the LL group than in the WW and WL groups if facial coloration has a function in emotional or social signaling.

### Tissue clearing and immunohistochemistry

Three bred discus fish *S. aequifasciatus* individuals (22.5–33 mm SL) were used for tissue clearing and immunohistochemistry, primarily following the methods described in Pende et al. (2020)^42^. The fish were deeply anesthetized with benzocaine, and their head and body were sectioned along a plane extending from the base of the pelvic fin to approximately the third to fifth spine of the dorsal fin. After removing the internal organs and creating small holes in the cornea, the heads were fixed in 4% (w/v) paraformaldehyde at 4°C overnight. The specimens were washed four times (30 minutes each) with PBS (pH 7.4) at room temperature and incubated in 0.2 M EDTA (pH 8.0) at 4°C for 48 hours, with the solution refreshed after 24 hours. After another four PBS washes at room temperature, they were transferred to pre-chilled acetone and incubated at 4°C for 1 hour. The acetone was then refreshed, and the specimens were incubated overnight at −20°C, followed by another refreshment and incubation overnight at room temperature. After being washed four times with PBS at room temperature, the specimens were treated with 3% (w/w) H_2_O_2_ in 0.8% (w/v) KOH for about 160 minutes at the same temperature under intense light exposure. Subsequently, they were washed with 1% (v/v) Triton X-100 in PBS (PBST) for two nights at 4°C and placed under vacuum for 60 minutes to remove gas bubbles. They were then incubated in Solution-1.1 (pH 11.5) for two nights at 37°C, with the solution being refreshed after the first night. This solution consisted of 10% (v/v) THEED (Sigma-Aldrich, 87600-100 ml), 5% (v/v) Triton X-100, and 5% (w/v) urea in dH_2_O, and its pH was adjusted to 11.5 using 30% (w/w) ammonia solution. After five PBST washes (10 minutes each), the specimens were treated with 1% (w/v) bovine serum albumin (BSA; Proliant Biologicals, PRL68700-25G) in PBST overnight at 4°C. Then, they were incubated in 1:1,000 primary antibody (rabbit anti-TH; Merck Millipore, AB152) in PBST with 0.5% BSA for 4 days at 4°C, followed by [*add number*] PBST washes for 2 days at 4°C. Next, the specimens were incubated in 1:200 biotinylated goat anti-rabbit IgG secondary antibodies (Vector Laboratories, BA-1000) in PBST with 0.5% BSA for 4 days at 4°C, washed in PBST for 2 days at 4°C, and incubated in a mixture of 1:100 avidin and 1:100 biotinylated horseradish peroxidase (Vector Laboratories, PK-6100) in PBST for 4 days at 4°C, followed by PBST washes for 2 days at 4°C. The specimens were stained in a reaction solution containing 0.05% (w/v) 3,3’-diaminobenzidine, 0.01% H_2_O_2_, 0.04% (w/v) nickel ammonium sulfate, 0.5% (v/v) Triton X-100, and 0.1 M phosphate buffer (pH 7.4) in ultra-pure water, followed by [*add number*] PBST washes for 1 day at room temperature. For refractive index matching, the three specimens were incubated in CUBIC-R+(N) (45% [w/w] antipyrine, 30% [w/w] nicotinamide, 0.5% [v/v] N-butyldiethanolamine in ultra-pure water) (Matsumoto et al. 2019) for 3 days at room temperature. The eye-patch regions were then observed and photographed under a microscope fitted with a digital camera (Olympus, SZX10 with DP25).

### Ex vivo administration of NA and ATP to scale melanophores

Three bred discus fish *S. aequifasciatus* individuals (21–24 mm SL) were anesthetized with benzocaine, and a total of 20 scales (ATP exposure group, n = 5; control for ATP exposure group, n = 5; NA exposure group, n = 5; control for NA exposure group, n = 5) were collected from the eye-patch region and kept in ice-cold PBS for a short period until the experiment was started.

The scales were placed on a concave slide glass filled with PBS, secured by pinching the edges with a polypropylene sheet, and observed under a microscope (Olympus, BX53). No melanophores were present in the pinched areas. It was difficult to completely standardize the physiological state of each individual at the time of scale collection and the neural innervation status across all scales. Therefore, the ATP exposure group was pre-treated with NA, and the NA exposure group was pre-treated with ATP, and changes in melanosome movement due to neurotransmitter exposure administration were specifically quantified. After removing the PBS from the concave slide, 0.1 mM ATP in PBS was added to the NA exposure group, and 0.1 µM NA in PBS was added to the ATP exposure group, followed by incubation at room temperature for 2 minutes. Subsequently, the solutions were removed, and the scales were washed by adding and quickly removing PBS. A total of 1 µM NA in PBS or PBS was added to the NA exposure group, and a total of 10 mM ATP in PBS or PBS was added to the ATP exposure group.

Considering the moment each solution was applied as 0 seconds, photographs were taken using a microscope fitted with a camera (Olympus, BX53 with DP74) at 1 second and every 10 seconds after that for 120 seconds. All images were adjusted for white balance using Adobe Photoshop (version 25.9.1) and analyzed using ImageJ (version 2.14.0.). To exclude xanthophores, the images were split into the three primary colors, and each monochrome one was used in the analysis. Subsequently, one well-captured melanophore (e.g., focus alignment) was randomly selected per scale, and its area was set as the ROI. Then, the area showing a grayscale above the identical threshold for all images within the ROI was calculated. The rate of variation in melanosome dispersion (%) was calculated using the following formula:

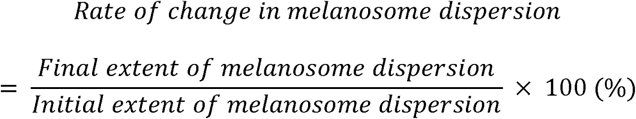

### Statistical analysis

All statistical analyses were performed in R (v. 4.3.3) ^77^ using LMMs or GLMMs in the lme4 package ^78^.

To evaluate whether facial coloration in subordinate individuals mitigated harassment by dominant individuals, the probability of continued harassment was analyzed as a binary variable within a binomial GLMM, with experimental pair ID incorporated as a random effect. The variation in facial coloration, specifically whether the facial pattern transitioned before or after the initial aggression by dominants (A to A, A to P, P to A, or P to P), was included as an explanatory variable. Given that changes in facial coloration were observed to significantly affect the likelihood of continuous harassment, post-hoc testing was conducted using separate binomial GLMMs, again treating experimental pair ID as a random effect. For these post-hoc GLMMs, the significance level (α) was adjusted via Bonferroni correction to α = 0.0083.

With regard to the experimental manipulation of facial coloration, a negative binomial GLMM was employed to analyze the data, with the experimental pair ID included as a random effect. This model examined the two-way interaction between the experimental groups (WW, WL, and LL pairs) and observation days (1, 3, 5, 7, and 9) as explanatory variables. Furthermore, the reduction in body weight during the experiment was analyzed using an LMM, again incorporating experimental pair ID as a random effect. The response variable was the percentage reduction in body weight (calculated as post-experiment body weight divided by pre-experiment body weight). The explanatory variables included the experimental group (WW, WL, and LL pairs), the sex of focal individuals, their status (light-treated or wild-type), and their social rank (dominant or subordinate).

Finally, the rate of variation in melanosome dispersion was examined upon exposure to ATP and NA, which act as sympathetic neurotransmitters. Prior to analysis, the black area in the selected melanophore (measured in pixels) was log-transformed (log10) to achieve normalization. An LMM was employed in this analysis, with individual ID included as a random effect. The two-way interaction between time and treatment (either ATP or PBS control) was modeled [*using?*] as an explanatory variable. An analogous analysis was conducted for the NA experiment. It should be noted that the results of these analyses are presented in Fig. 5 as the percentage change in melanosome dispersion relative to the baseline measurements at the start of the experiment for visualization.

## Acknowledgments

We thank the members of the Animal Sociology Laboratory of Osaka City University and the Kutsukake Research Group of Graduate University for Advanced Studies for their helpful comments. This study was supported financially by HAKUBI Project and KAKENHI (nos. N19K23765, N19KK0189, 20J01170, and 23KK0131 to S.S; nos. 23H03868 to S.S. and K.F.).

## Author contribution

S.S., K.F., A.K., S.T., M.K., and N.K. designed the study. S.S., K.F., S.M., T.B., and K.K. collected the data. S.S. and K.F. conducted the analyses. S.S., H.M., K.K., and K.F. wrote the manuscript with input from all the authors.

## Ethics statement

All the experimental protocols were approved by the Animal Care and Use Committees at Osaka City University and Kitazato University and adhered to the ASAB/ABS guidelines for the treatment of animals in behavioral research.

## Data availability

The behavioral data that support the findings of this study are available in Dryad XXX.

## Supplementary

**Supplemental information S1** Detailed observation methods for face and body coloration change for discus fish *Symphysodon aequifasciatus* and four species from tribe Heroini (Banded cichlid *Heros severus*, triangle cichlid *Uaru amphiacanthoides*, flag cichlid *Mesonauta festivus*, and angelfish *Pterophyllum scalare*).

**Supplemental movie S2** Typical example of facial coloration transition in the discus fish *Symphysodon aequifasciatus*. To facilitate the viewing of changes in facial coloration patterns, we presented a model of the reproductive partner on the monitor and observed its behavior based on the method described in Satoh et al. (2016) ^25^.

**Supplemental Figure S3.** Coloration associated with social behaviors in light-treated *S. aequifasciatus*. The bars represent the possibility of coloration patterns emerging when individuals engage in specific social behaviors. Three identifiable coloration patterns were detected: 1) absence of any vertical black bar composed of melanophores; 2) presence of only one bar consisting of melanophores, eye-patch pattern, and 3) presence of multiple bars throughout the body. These observations were made in the same manner as in *behavioral observation I* (see Methods).

