## supplementary S3 for "Social benefits of facial experssion in a cichlid fish: Testing the face concentration hypothesis"

**Supplemental movie S1** Typical example of facial coloration transition in the discus fish *Symphysodon aequifasciatus*. To facilitate the viewing of changes in facial coloration patterns, we presented a model of the reproductive partner on the monitor and observed its behavior based on the method described in Satoh et al. (2016) ^25^.

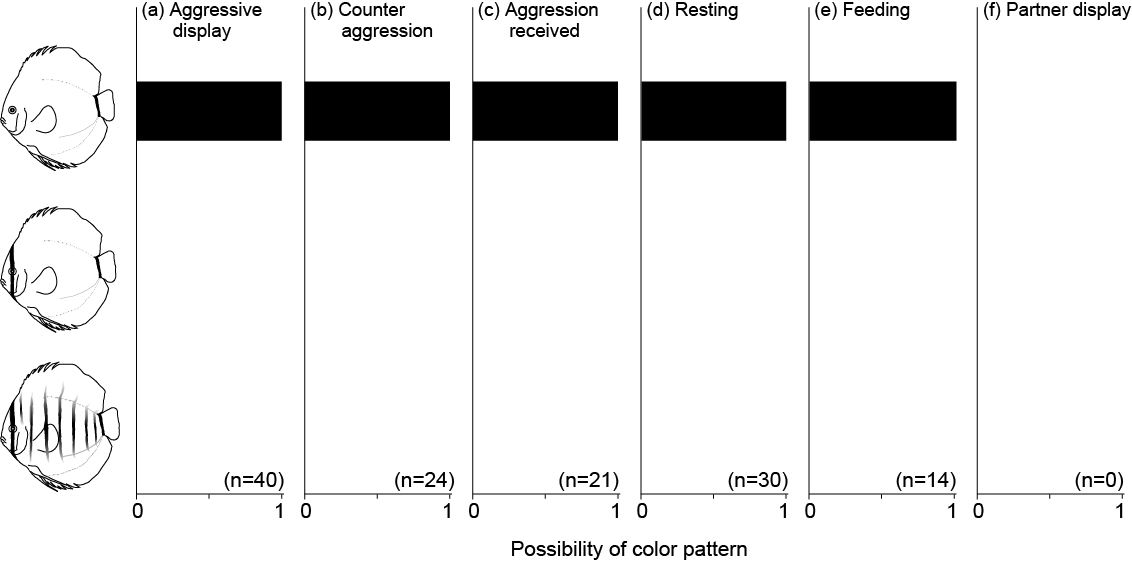

**Supplemental Figure S2** Coloration associated with social behaviors in light-treated *S. aequifasciatus*. The bars represent the possibility of coloration patterns emerging when individuals engage in specific social behaviors. Three identifiable coloration patterns were detected: 1) absence of any vertical black bar composed of melanophores; 2) presence of only one bar consisting of melanophores, eye-patch pattern, and 3) presence of multiple bars throughout the body. These observations were made in the same manner as in *behavioral observation I* (see Methods).
