## supplementary S1 for "Social benefits of facial experssion in a cichlid fish: Testing the face concentration hypothesis"

Ten adult fish, obtained from ornamental fish suppliers, were maintained by species in a 360 L tank (length × width × height = 120 × 60 × 50 cm). The sexes of these individuals were unknown, but they were at or near body size sexually matured. During rearing, experimental individuals were fed mosquito larvae three times daily, and they exhibited social interactions in the tanks. After a habituation period of more than one week, we examined the relationship between their body patterns and aggressive interactions. A camera (OMD-2, Olympus Corporation, Tokyo, Japan) was used to capture moments when the focal individuals displayed aggression toward conspecifics and when they were attacked. If the focal individual exhibited counter-aggression or lateral display when attacked against aggressor, it was not considered attacked and was not photographed. Filming was restricted to instances in which the individuals were oriented parallel to the camera whenever possible, and subsequent filming took place within one hour of the initial filming (when they exhibited aggression or were attacked by a tank mate).

Some individuals neither exhibited aggression nor experienced aggression from tank mates because of a fixed dominance hierarchy. Individuals that exhibited only one of these behaviors during the one-hour behavioral observation and filming were removed from the observation tank using a hand net and replaced with individuals from the stock tank (121.5 L or 182.3 L). After an acclimation period of at least one week, filming was repeated for the replaced individuals. This observation was conducted for discus fish (*Symphysodon aequifasciatus*, n = 10) and four related species: banded cichlid (*Heros severus*, n = 10), triangle cichlid (*Uaru amphiacanthoides*, n = 10), flag cichlid (*Mesonauta festivus*, n = 10), and angelfish (*Pterophyllum scalare*, n = 10).

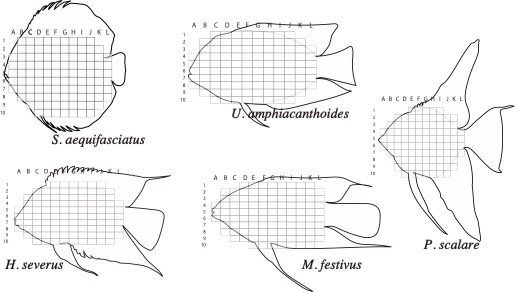

Because it was challenging to establish a color correction standard near the moving focal individuals within the tank, we visually quantified color changes for each aggressive interaction. The captured images were printed on glossy paper and presented to the raters in individual combinations. Concurrently, the raters were provided with illustrations depicting only the target species’ outlines and asked to mark, within a grid system, the areas that exhibited different coloration between the two images. The illustrations were divided into 12 × 10 grids, totaling 96 to 107 grids depending on the body shape of focal species. Note that the color change was assessed under blind conditions; the raters were unaware of our hypotheses or research concepts. These data were compiled to generate a heat map of color change for each individual (see below).

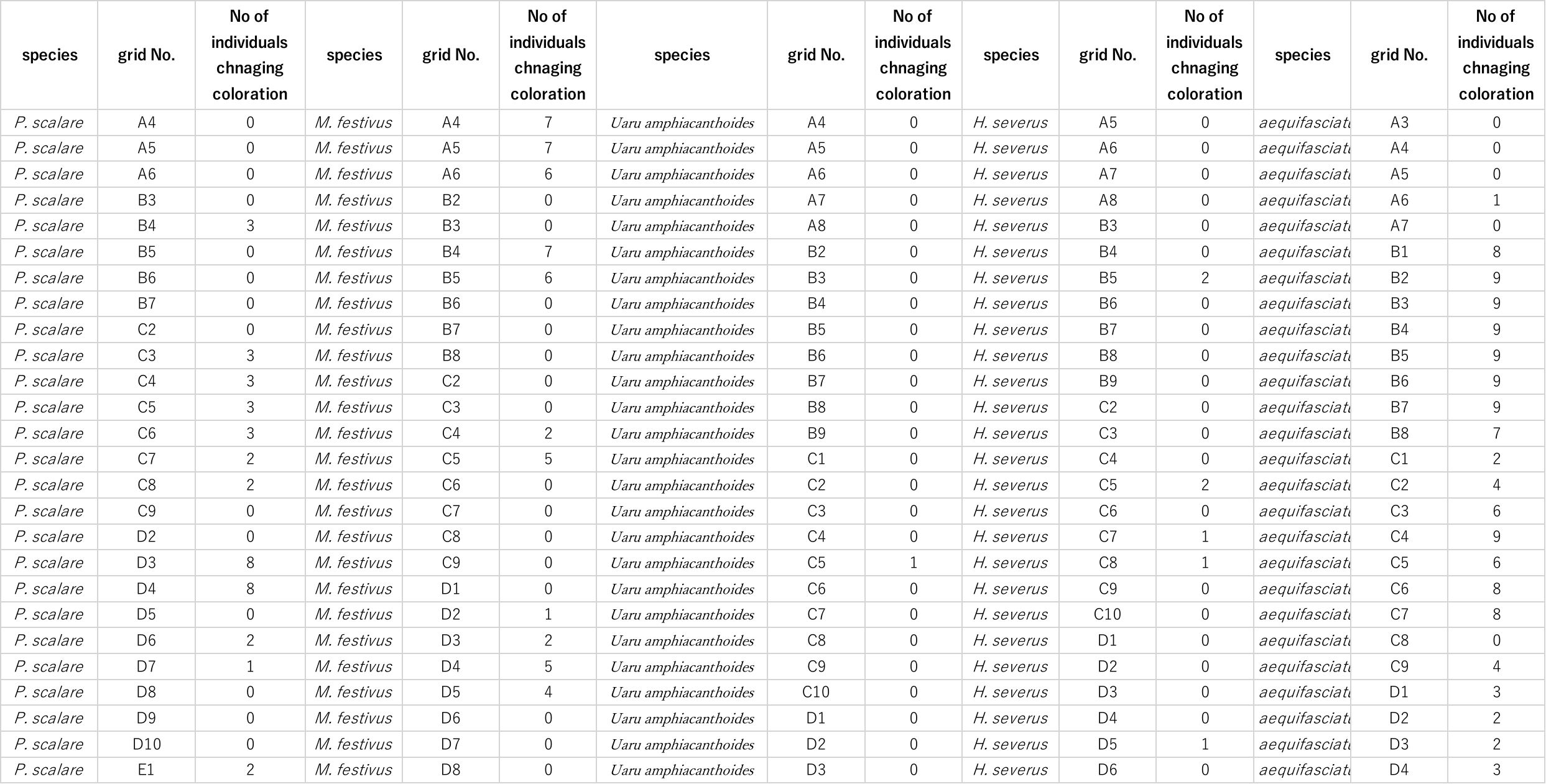

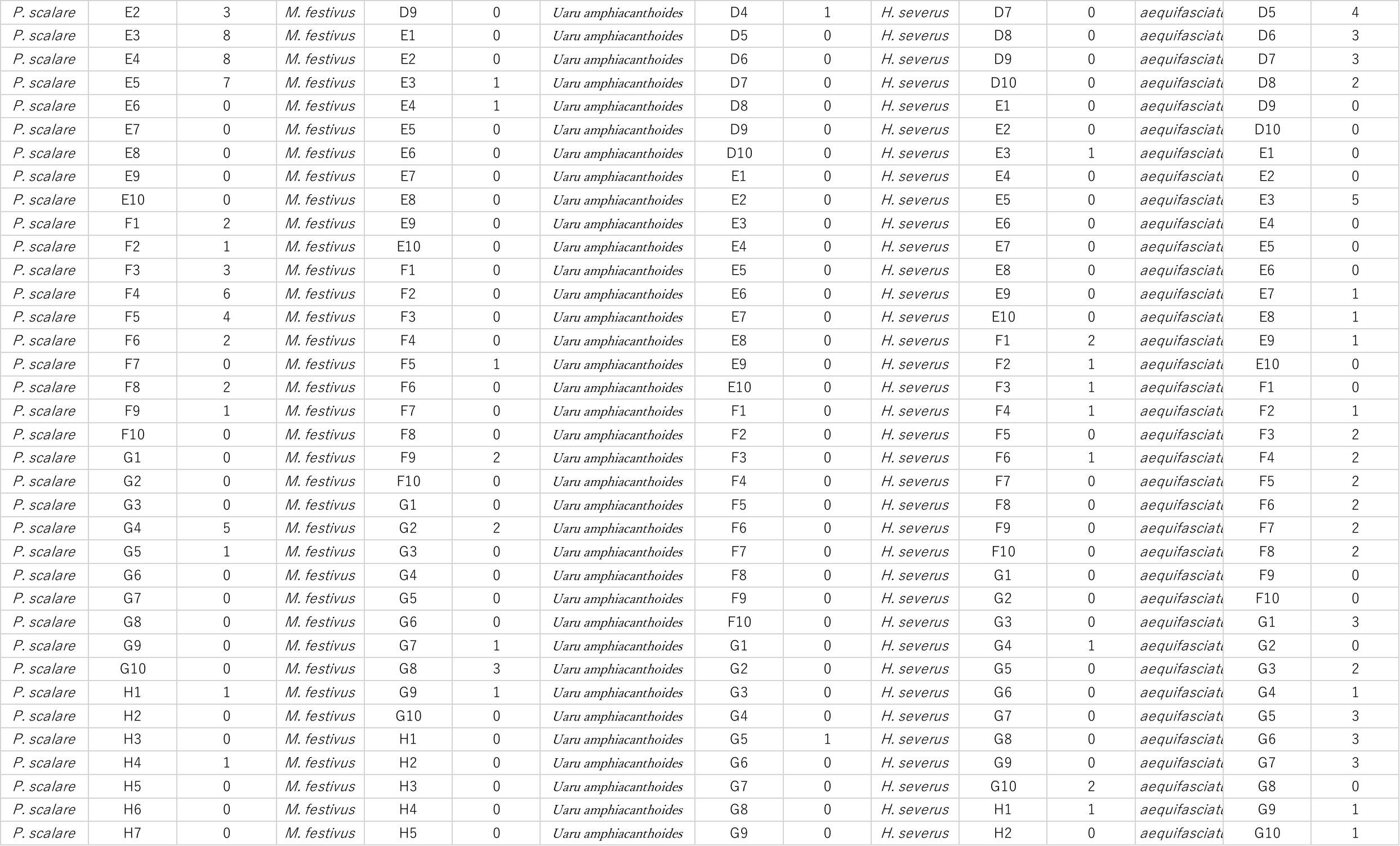

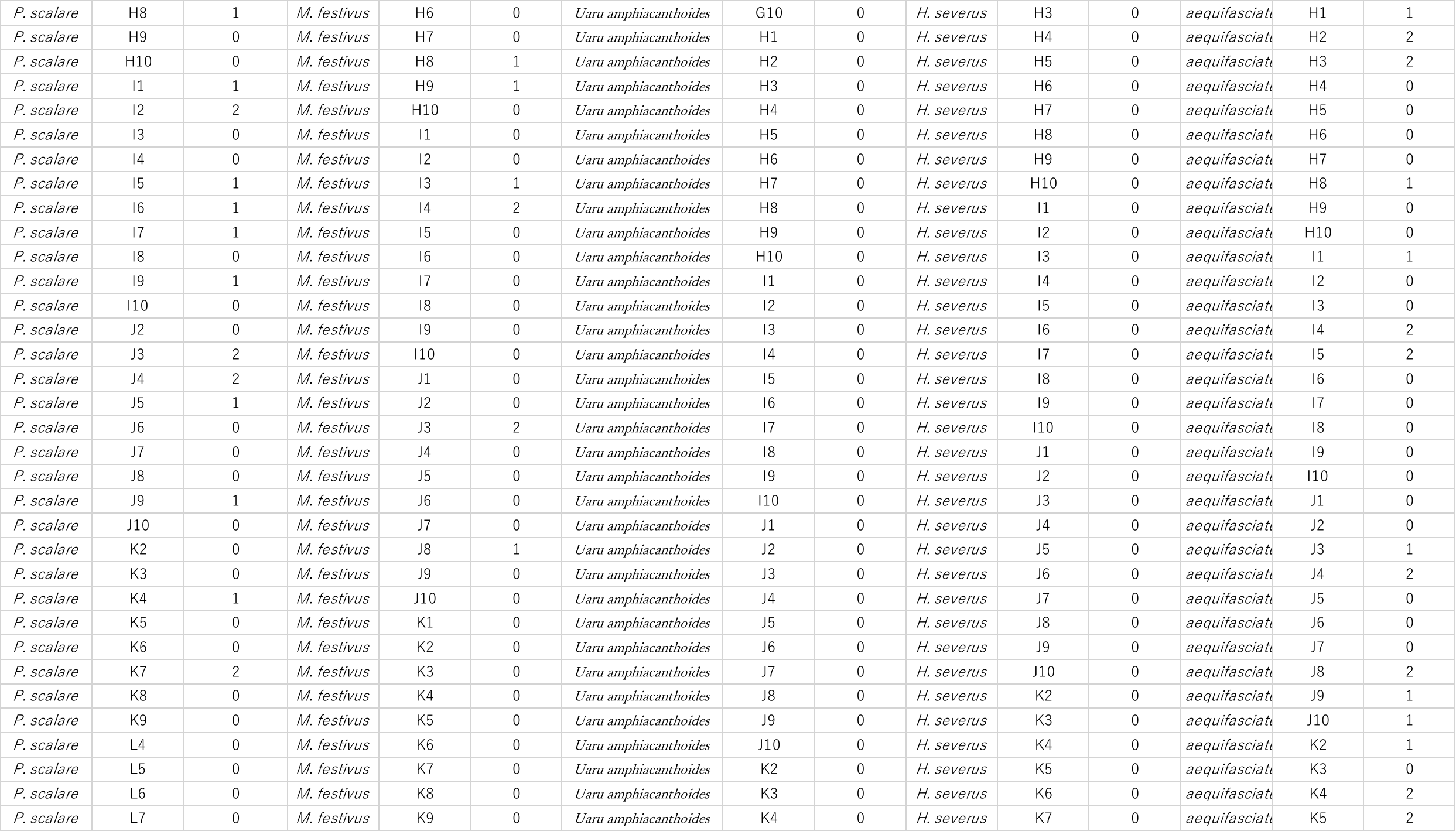

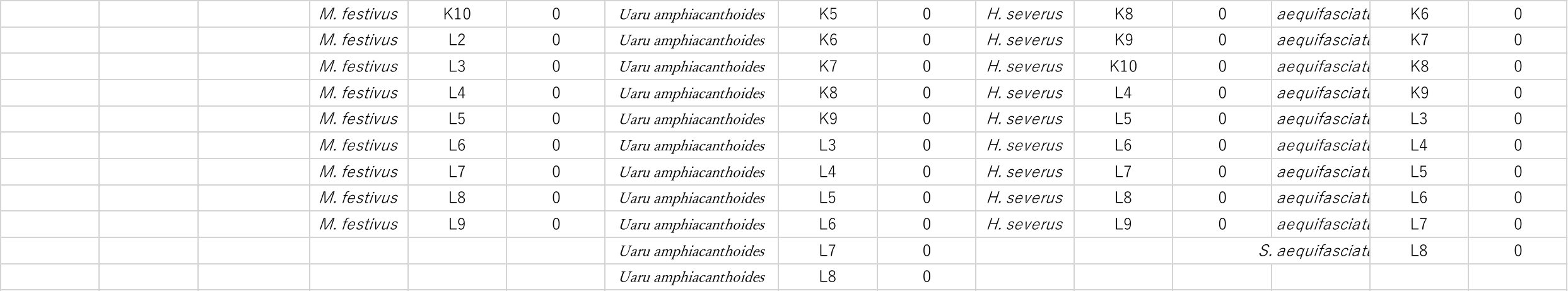
